# TRiC-assisted folding of class I HDAC family proteins regulated by distinct co-chaperone and cofactor networks

**DOI:** 10.1101/2025.05.21.655272

**Authors:** Zuyang Li, Qiaoyu Zhao, Wanying Jiang, Qianqian Song, Xuehai Zhou, Xiangyi Shi, Qing Zhang, Yanxing Wang, Yinghong Lin, Yue Yin, Chen Pan, Yao Cong

## Abstract

Class I histone deacetylases (HDACs), including HDAC1, HDAC2, HDAC3, and HDAC8, are essential for diverse cellular processes. Although the chaperonin TRiC is implicated in the activation of class I HDACs, the underlying mechanisms remain elusive. Using cryo-electron microscopy (cryo-EM), cross-linking mass spectrometry (XL-MS), and biochemistry analyses, we established class I HDACs as novel TRiC substrates and elucidate TRiC-assisted folding of HDAC1 and HDAC3 during its ATPase cycle, orchestrated by distinct co-chaperone and cofactor networks. In the closed TRiC chamber, both HDAC1 and HDAC3 adopt near-native states with shared binding interfaces. However, their open-state configurations diverge: Hsp70 and PDCD5 engage atop and within TRiC, respectively, for HDAC3, whereas prefoldin (PFD) binds atop TRiC for HDAC1, suggesting roles in substrate delivery and folding modulation. Furthermore, an unexpected bent conformation of CCT4, detected in TRiC-HDAC1 complex, may facilitate co-chaperone dissociation from TRiC. In contrast, HDAC8 folds independently of TRiC. Our study reveals the mechanism governing TRiC-assisted folding of class I HDACs in orchestration of dynamic co-chaperone/cofactor network, shielding new lights on the sophisticated regulatory landscape of TRiC, and open promising avenues for designing peptides or small molecules to selectively modulate TRiC-assisted folding of class I HDACs and other substrates.

## Introduction

The eukaryotic group II chaperonin TRiC (also known as CCT) is indispensable for folding approximately 10% of cytosolic proteins, including key cytoskeletal components (actin and tubulin), the cell cycle regulator CDC20, and the VHL tumor suppressor^1–3^. TRiC facilitates substrates folding through an ATP-driven conformational cycle^4–7^. This large, hetero-oligomeric complex (∼1 MDa) consists of two stacked octameric rings, each formed by eight distinct subunits (CCT1–CCT8)^8–10^. TRiC works in concert with a suite of molecular chaperones—including Hsp70^11,12^, prefoldin (PFD)^13,14^, phosducin-like proteins (PhLPs)^15–17,22^, and programmed cell death protein 5 (PDCD5)^18,19^—to orchestrate protein folding pathways. Hsp70 stabilizes unfolded or partially folded proteins during the initial stages of folding^20^. PFD, a jellyfish-shaped hetero-oligomeric complex, binds the open state TRiC and dissociates upon ATP-driven lid closure^13,14^. Hsp70 and PFD may direct the transfer of certain substrates to TRiC^11,12,14^. PhLPs (including PhLP1, PhLP2, and PhLP3) modulate TRiC-mediated protein folding^21^. PDCD5, which associates with nearly all open-state TRiC in HEK293 cells^19^, is proposed to function as a modulator of TRiC, potentially related to β-tubulin folding^18^. Over the past decade, sustained research efforts have clarified the roles of PFD and PhLPs in TRiC-mediated folding of tubulin and actin^13,14,14,15,22^. However, the intricate mechanisms by which Hsp70, PFD, and PDCD5 cooperate with TRiC to fold a broader range of substrates remain not fully elucidated.

Histone deacetylases (HDACs) are a superfamily of chromatin-modifying enzymes that regulate gene expression by catalyzing the removal of acetyl groups from histones^23^. This deacetylation induces chromatin condensation, reduces DNA accessibility, and suppresses gene transcription^24^. Conversely, HDAC inhibition induces histone hyperacetylation and transcriptional activation, a mechanism exploited for therapeutic strategies against cancer and other epigenetic disorders. The human HDAC family is categorized into four classes based on sequence homology^23,25^. Class I HDACs—including HDAC1, HDAC2, HDAC3, and HDAC8—are particularly significant due to their ubiquitous expression and involvement in critical cellular processes, including cell cycle regulation, differentiation, and development^26^.

HDAC1, HDAC2, and HDAC3 predominantly localize in the nucleus, functioning as core components of large multiprotein complexes^27^. While HDAC1 and HDAC2 typically function within multiprotein co-repressor complexes such as NuRD^28^, CoREST^29^, and Sin3A^30–32^, HDAC3 is uniquely associated with the SMRT/NCoR co-repressor complex^33^. In contrast, HDAC8 is the only class I HDAC exhibiting full enzymatic activity independently of large complexes^34,35^. Dysregulation of class I HDACs is implicated in numerous pathologies, including cancer, neurodegenerative diseases, and inflammatory disorders, rendering them as promising therapeutic targets^26^. In vitro, HDAC3 gains its histone-deacetylation activity through stable association with the conserved deacetylase-activating domain (DAD) of NCOR1 and SMRT^36,37^—an energy-dependent process facilitated by the TRiC complex^38^. However, the precise mechanism by which TRiC facilitates HDAC3 activation remains elusive, and whether other class I HDACs similarly depend on TRiC for activation is unknown.

To address these questions, we purified HDAC1, HDAC3, and HDAC8, along with their endogenously interacting proteins, from HEK293F cells. Employing cryo-electron microscopy (cryo-EM), cross-linking and mass spectrometry (XL-MS), and biochemical and functional assays, we identified a network of chaperones interacting with HDAC3 and HDAC1. We elucidated distinct TRiC-assisted folding mechanisms for HDAC3 and HDAC1, mediated by specific co-chaperone and cofactor networks: Hsp70 and PDCD5 for HDAC3, and PFD for HDAC1. In contrast, HDAC8 folds independently of TRiC. Notably, parallel functional co-chaperones or cofactors associate with TRiC in a mutually exclusive manner, revealing a finely tuned specificity in their interactions. This study highlights the complexity of class I HDAC folding, assembly, and activation, regulated by the precise coordination of TRiC with its co-chaperone and cofactor networks, and underscores the specificity of these networks for each individual class I HDAC.

## Results

### HDAC3 folding is mediated by TRiC and potentially involves Hsp70

To characterize the HDAC3 protein interactome, we expressed Flag-tagged HDAC3 in HEK293F cells and affinity-purified it along with its endogenously interacting partners. Mass spectrometry (MS) analysis of the HDAC3 immunoprecipitate revealed that the predominant interacting proteins were molecular chaperones, including Hsp70 variants (with HSPA1B as the predominant isoform), all eight subunits of the TRiC complex, and Hsp90 (Fig. 1A, Fig. S1A). Using glycerol density gradient centrifugation, we purified two distinct complexes (Fig. 1B). MS analysis of fractions 13-14 identified a complex containing Hsp90, Hsp70, HOP, and HDAC3 (Fig. 1C, Fig. S1B), resembling the client-loading complex of the glucocorticoid receptor^39^. This suggests a potential role for Hsp90 in HDAC3 regulation, though the nature of their interaction warrants further investigation. The other complex, from fractions 19-20, comprises all eight TRiC subunits, Hsp70, and HDAC3, and is the focus of this study (Fig. 1D, Fig. S1C). Native electrophoresis and immunoblotting confirmed the co-presence of HDAC3 and Hsp70 together with TRiC (Fig. 1E, Fig. S1D). Furthermore, XL-MS analysis detected direct interactions between HDAC3 and TRiC, as well as between Hsp70 and TRiC (Fig. 1F, S1E, Table S2). Together, these data establish the formation of a stable complex involving TRiC, HDAC3, and Hsp70.

**Fig. 1.**
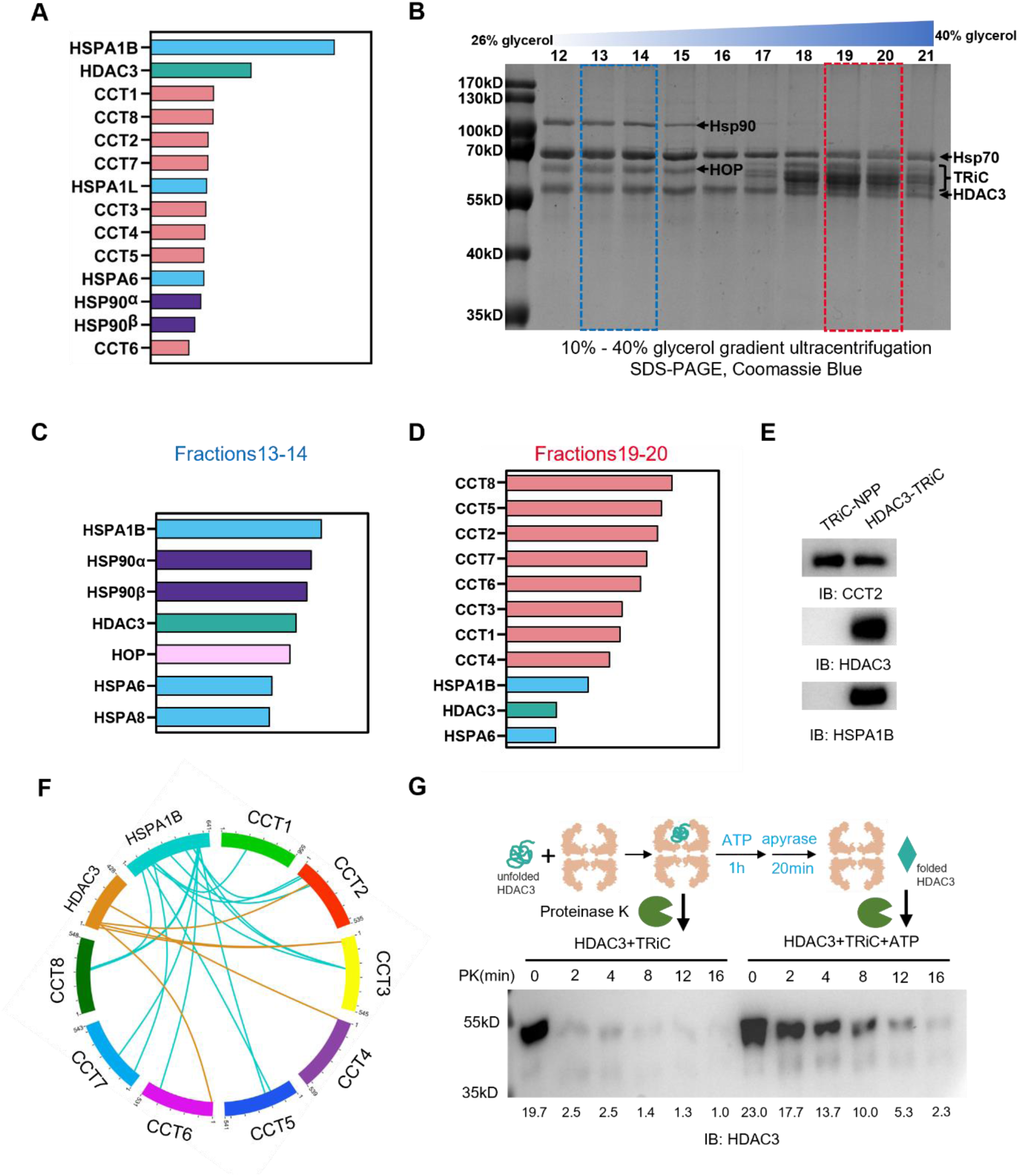
A chaperone network interacting with HDAC3. (A) Mass spectrometry (MS) identified top 14 HDAC3-interacting proteins following immunoprecipitation of Flag-tagged HDAC3, ranked by peptide-spectrum matches (PSM). (B) Proteins immunoprecipitated with Flag-tagged HDAC3 were analyzed by 10%-40% glycerol gradient ultra-centrifugation. Fractions, collected from top to bottom, were analyzed by SDS-PAGE and Coomassie Brilliant Blue staining. (C) MS identified top 7 chaperones in fractions 13-14. (D) MS identified top 11 chaperones in fractions 19-20. (E) Proteins extracted from the native gel were analyzed by western blotting, detecting HDAC3, CCT2, and HSPA1B. (F) XL-MS analysis of the HDAC3-TRiC complex revealed cross-links between HDAC3, Hsp70, and TRiC, filtered using the best E-value (1.00E−02) and spectral counts of at least 2 as the threshold to exclude low-confidence data. (G) In the Proteinase K sensitivity assay, denatured HDAC3 (purified from HEK293F cells) was incubated with TRiC in the presence or absence of ATP. After the reaction, residual ATP was hydrolyzed by apyrase, and HDAC3 was detected by western blotting.

To determine whether HDAC3 is a TRiC substrate, we performed a proteinase K (PK) resistant assay, which distinguishes unstructured polypeptides—susceptible to degradation by low concentration of the non-specific protease PK—from folded domains that exhibit increased PK resistance^14,40,41^. Purified TRiC and denatured HDAC3 (Fig. S1F-G) were incubated together, and ATP was added to induce TRiC ring closure^42^. This delayed HDAC3 degradation by PK (Fig. S1H). Strikingly, this protective effect persisted even after adding apyrase (Fig. 1G), which catalyzes ATP hydrolysis^43^, triggering TRiC ring reopening and substrate release. These findings suggest that HDAC3’s PK resistance arises from TRiC-mediated folding rather than ATP-induced TRiC closure. Taken together, our results suggest that HDAC3 is a substrate of TRiC, with Hsp70 potentially involved in its folding.

### Hsp70 interacts with TRiC to facilitate TRiC-mediated HDAC3 folding

To elucidate the role of Hsp70 in TRiC-mediated folding of HDAC3, we conducted cryo-EM analysis of the TRiC-HDAC3-Hsp70 complex without added nucleotides. This yielded two cryo-EM maps (Fig. S2A): one depicting an empty TRiC in an open conformation (3.72 Å resolution), and another revealing TRiC, also in an open state, with additional density atop one ring (termed the S1 state, 3.95 Å resolution; Fig. S2B-D). In the S1 state, the TRiC conformation resembles the nucleotide partially preloaded (NPP) state of human TRiC^44^, with the extra density atop TRiC primarily associated with the CCT4 A-domain (Fig. 2A-B and E). Both maps show a central density between the two equators (Fig. S2E), consistent with prior cryo-EM studies of open TRiC from human, bovine, and yeast, attributed to its unstructured N- and C-terminal tails^4,7,10,15,44,45^.

**Fig. 2.**
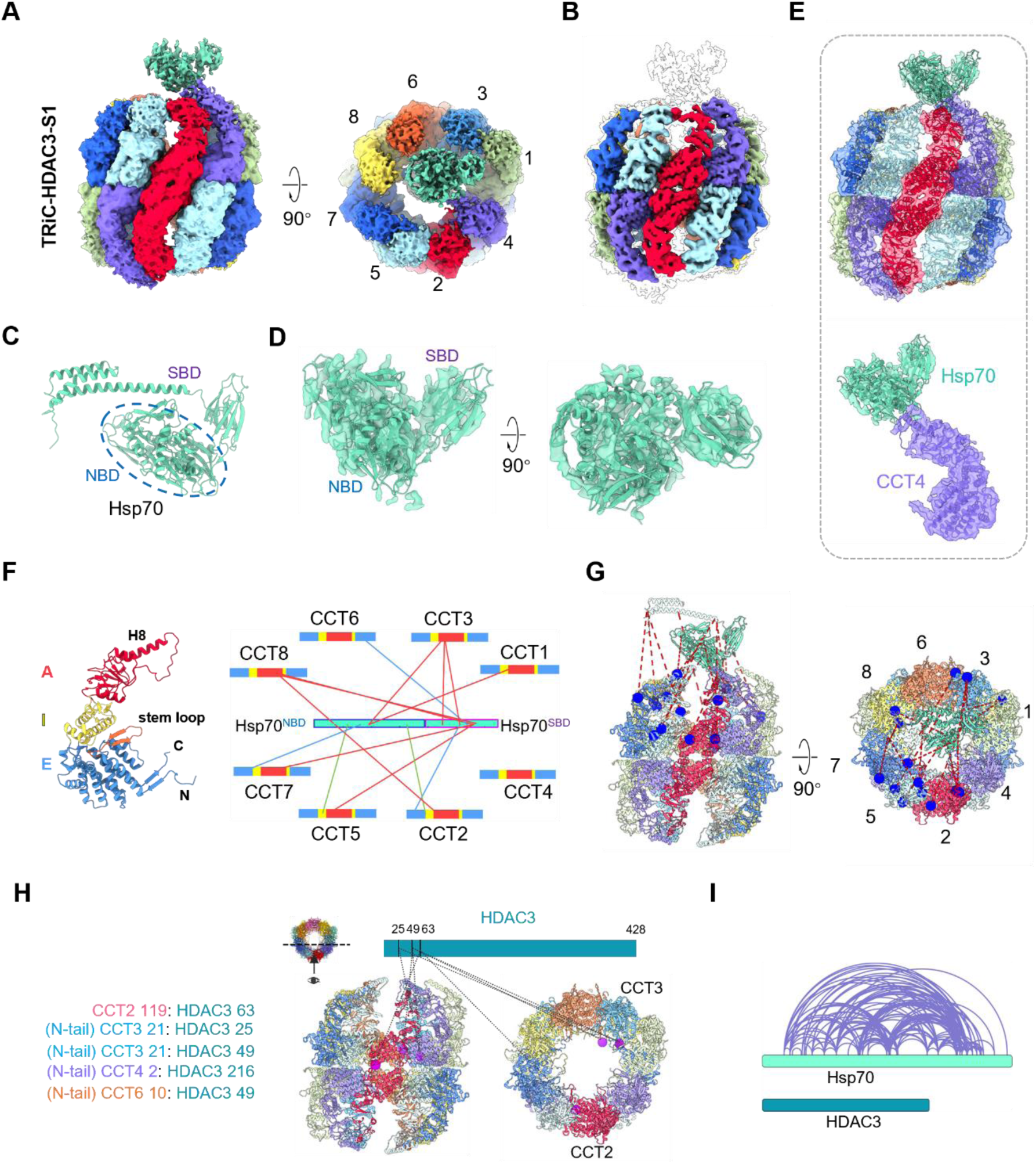
Cryo-EM structures of HDAC3-TRiC in the NPP state. (A) Different views of the unsharpened cryo-EM map of HDAC3-TRiC-S1 state, exhibiting an extra density (light green) atop TRiC. TRiC subunits are distinctly colored, with this color scheme maintained in subsequent figures. (B) The sharpened HDAC3-TRiC-S1 map (with color) within the the unsharpened map (transparent envelope). (C) AlphaFold2-predicted Hsp70 model, containing the nucleotide-binding domain (NBD, blue circle) and substrate-binding domain (SBD). (D) Visualization of the extra density fits well with the Hsp70 model. (E) Map-model fitting of the S1 state, and a close-up of the extra density contacting CCT4. (F) XL-MS cross-links between Hsp70 and TRiC, mapped onto their linear bar diagrams, predominantly localized in the TRiC A-domain (in red). (G) Intermolecular Hsp70-TRiC cross-links mapped onto the structural model. (H) Intermolecular HDAC3-TRiC cross-links mapped onto the TRiC model. Cross-links involving disordered TRiC N-/C-terminal tails, not depicted in the model, are listed on the left. (I) Self cross-links of Hsp70 and HDAC3 in fractions 7-8 from glycerol density gradient centrifugation.

Interestingly, the extra density atop the TRiC ring resembles Hsp70 in size and shape, though it lacks the C-terminal three α-helixes (Fig. 2C-D). Its relatively weak appearance reflects intrinsic dynamics and compositionally heterogeneity. Our XL-MS analysis further identified 15 crosslinks between Hsp70 and all eight TRiC subunits, predominantly to the A- and I-domains of TRiC on its outer surface (Fig. 2F-G), corroborating our structural model and indicating that this density indeed represents dynamically bound Hsp70. Additionally, five HDAC3-TRiC crosslinks were detected, mapped primarily within the TRiC cavity, specifically to the E-domains of CCT2, CCT3, CCT4, and CCT6 (Fig. 2H). Taken together, our cryo-EM and XL-MS data indicate that Hsp70 docks atop TRiC—distinct from a previously reported site outside TRiC near CCT2 subunit^12^—while HDAC3 resides within TRiC cavity.

Given that Hsp70 co-purifies endogenously with HDAC3 (Fig. 1B), its binding atop TRiC, and previous biochemical evidence that the folding of some substrates requires Hsp70-TRiC cooperation—where Hsp70 promotes substrate binding to TRiC^11,46^—we propose that Hsp70 may faciliate HDAC3 delivery to TRiC. Indeed, additional XL-MS analysis of glycerol gradient fractions 7-8 revealed abundant Hsp70 self-crosslinks, indicative of a folded state, whereas HDAC3 lacked such self-crosslinks (Fig. 2I). This pattern persisted for the complex in fractions 19-20 used for cryo-EM (Fig. S1E), suggesting that Hsp70 remains folded while HDAC3 is largely unstructured and dynamic. These findings reinforce the hypothesis that Hsp70 facilitates the delivery of unstructured HDAC3 to TRiC for folding.

### ATP-driven TRiC ring closure drives HDAC3 folding

To investigate whether TRiC assists in HDAC3 folding through its conformational cycle and to elucidate the mechanism, we performed cryo-EM analysis of the TRiC-HDAC3 complex in the presence of ATP-AlFx, an ATP-hydrolysis transition-state analog known to induce TRiC ring closure^5,15,44^. This revealed two TRiC conformational states: a closed state (36% of the population) and an open state (64%, discussed later; Fig. S3). For the closed TRiC, we applied symmetry expansion and focused 3D variability analysis (3DVA)^47^, yielding a 3.30 Å resolution map, termed TRiC-HDAC3-S2 (Fig. 3A, Fig. S4A-D). Notably, this map disclosed an additional density within the TRiC chamber (Fig. 3B), closely matching the HDAC3 model and exhibiting high-resolution features (Fig. 3C-D). Indeed, this density corresponds to HDAC3 in a nearly fully folded state (Fig. 3C), demonstrating that HDAC3 is a TRiC substrate, whose folding is facilitated by ATP-driven TRiC ring closure.

**Fig. 3.**
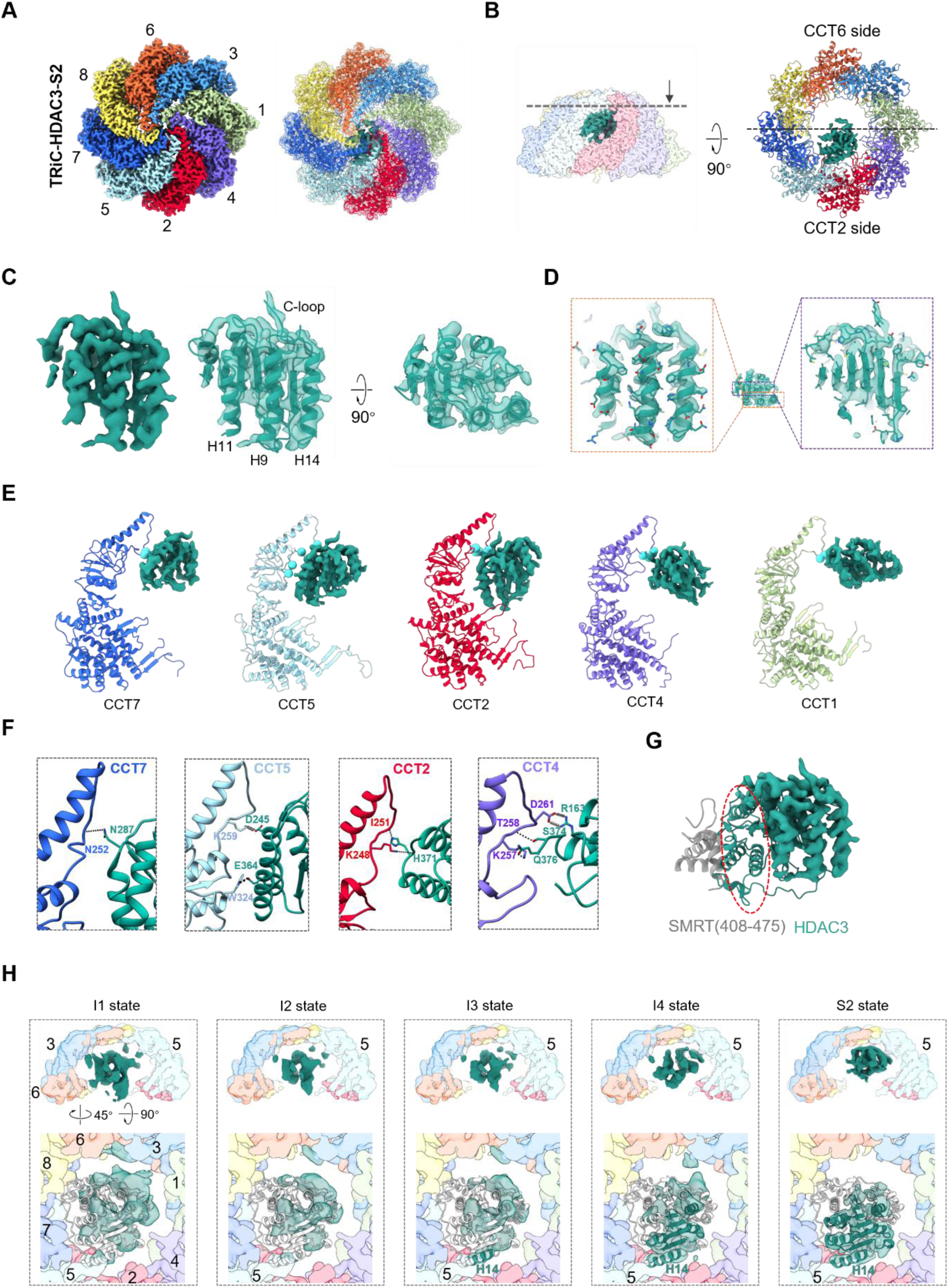
Cryo-EM structures of HDAC3-TRiC in the closed state. (A) Cryo-EM map and structure of the closed HDAC3-TRiC-S2 state, exhibiting an extra density (seagrass green) within the TRiC chamber. (B) Visualization of the extra density in the TRiC chamber, which fits well with the HDAC3 model (PDB: 4A69). (C) High-resolution structural features of HDAC3. (D) TRiC-HDAC3 interaction interface analysis, showing HDAC3 contacts with CCT7/5/2/4/1 subunits. TRiC residues close to HDAC3 (within 4 Å) are shown as cyan balls. (E) Detailed interaction networks between HDAC3 and TRiC, with hydrogen bonds as dashed lines and salt bridges as springs (style maintained in subsequent figures). (F) Fitting of the HDAC3 density with the HDAC3-SMRT-DAD complex (PDB: 4A69). The region of HDAC3 interacting with SMRT appears poorly folded (red circle). (G) Side and cut-away top views of HDAC3-TRiC maps in I1–I4 and S2 states, illustrating TRiC-assisted HDAC3 folding process. I1–I4 maps were derived using 3DVA; the S2 map is unsharpened one. Unfolded HDAC3 regions are shown in light grey, folded regions in cyan (bottom row).

Notably, HDAC3 engages with TRiC on the CCT2 side (Fig. 3B), similar to Gβ_5_^16^ but distinct from tubulin or actin, which bind on the CCT6 side^14,15,17,22,44^. Positioned beneath the TRiC dome, HDAC3 interacts with CCT7/5/2/4/1 subunits through its H11/13/16-helices and c-loop through H-bonds and salt bridges (Fig. 3E-F, Table S3). Although HDAC3 is nearly fully folded within TRiC chamber, the region corresponding to its interaction with the DAD of SMRT remains disordered (Fig. 3G). This suggests intrinsic flexibility in this interface, which likely becomes stabilized after being released from TRiC and upon binding to SMRT. These findings indicate that TRiC not only folds HDAC3 but may also play a role in assembling the HDAC3-SMRT complex.

### Cryo-EM reveals HDAC3 folding process within TRiC and the mechanism

To further capture the folding process of HDAC3 within TRiC, we performed 3DVA on the same closed-state cryo-EM dataset (Fig. S3). This analysis revealed four additional intermediate folding states of HDAC3, designated I1 to I4, alongside the folded TRiC-HDAC3-S2 state (Fig. 3H). Together, these five states provide a more comprehensive view of the HDAC3 folding process within the TRiC chamber. In the I1 state, the substrate density is less compact, lacking discernible structural features of HDAC3, and appears as a fluffy mass that contacts the TRiC chamber at multiple sites: two on CCT3 and one each on CCT5 and CCT7. As folding progresses from I2 to I4 states, the substrate density reduces its interactions to only CCT3 and CCT5, and becomes more structured. Notably, in the I3 state, the C-terminal Η14-helix emerges as the first discernible feature of HDAC3 (Fig. 3H). By the I4 state, additional structural elements surrounding Η14, including Η8, Η9, Η11, and strands S4-S6, become discernible. Finally, in the S2 state, the HDAC3 density appears compact and fully folded, with interactions now restricted to CCT5. Throughout this folding process, the substrate density shifts upward toward the TRiC dome and laterally toward the CCT2 side.

Collectively, these structural observations suggest a stepwise mechanism for TRiC-assisted HDAC3 folding. In the early I1 and I2 states, HDAC3 adopts a loosely assembled conformation, resembling its overall shape but appearing larger in size, and interacts with TRiC inner wall at multiple sites. In the I3 state, the H14 helix emerges as a pivotal interaction hub with the CCT5 subunit, potentially serving as a folding nucleus, facilitating the propagation of folding of adjacent regions. Finally, in the I4 and S2 states, folding propagates from H14 to nearby elements, followed by the remaining regions, which rapidly overcome energy barriers on the free-energy landscape, approaching the global minimum of the HDAC3 native state.

### Identification of PDCD5 in open TRiC chamber in the ATP-bound state

In the open-state TRiC map derived also from the TRiC-HDAC3-ATP-AlFx dataset (Fig. S3), representing the ATP-bound state^15,44^, an additional density appeared deep within the TRiC chamber, bound to E-domains CCT3/1/4 (Fig. 4A-B), while the atop Hsp70 density is absent. Focused processing produced a 3.32 Å resolution consensus map and a 3.36 Å local map (Fig. S5A-E), merged as TRiC-HDAC3-S3 for analysis (Fig. 4A). Distinct from HDAC3, this density suggests a unique cofactor bound to open TRiC in the presence of HDAC3 and nucleotide. Fitting AlphaFold2 ^48^ predicted models of MS-detected proteins (Fig. S1C) into the density identified PDCD5 as the best match (correlation coefficient of 0.88, Fig. 4C, S5F). PDCD5, a highly conserved protein upregulated during apoptosis^49^, has been proposed to modulate TRiC’s β-tubulin folding^18^. XL-MS detected extensive PDCD5-TRiC cross-links (Fig. 4D-E, Table S4), confirming this density as PDCD5. Notably, its position differs from a prior report suggesting PDCD5 binds the A-domain of CCT2^18^, but aligns with a recent cryo-ET study of *in situ* TRiC from HEK293 cells^19^. Further focused classification revealed TRiC binds one (56%) or two PDCD5 molecules (44%) in its chamber (Fig. S3).

**Fig. 4.**
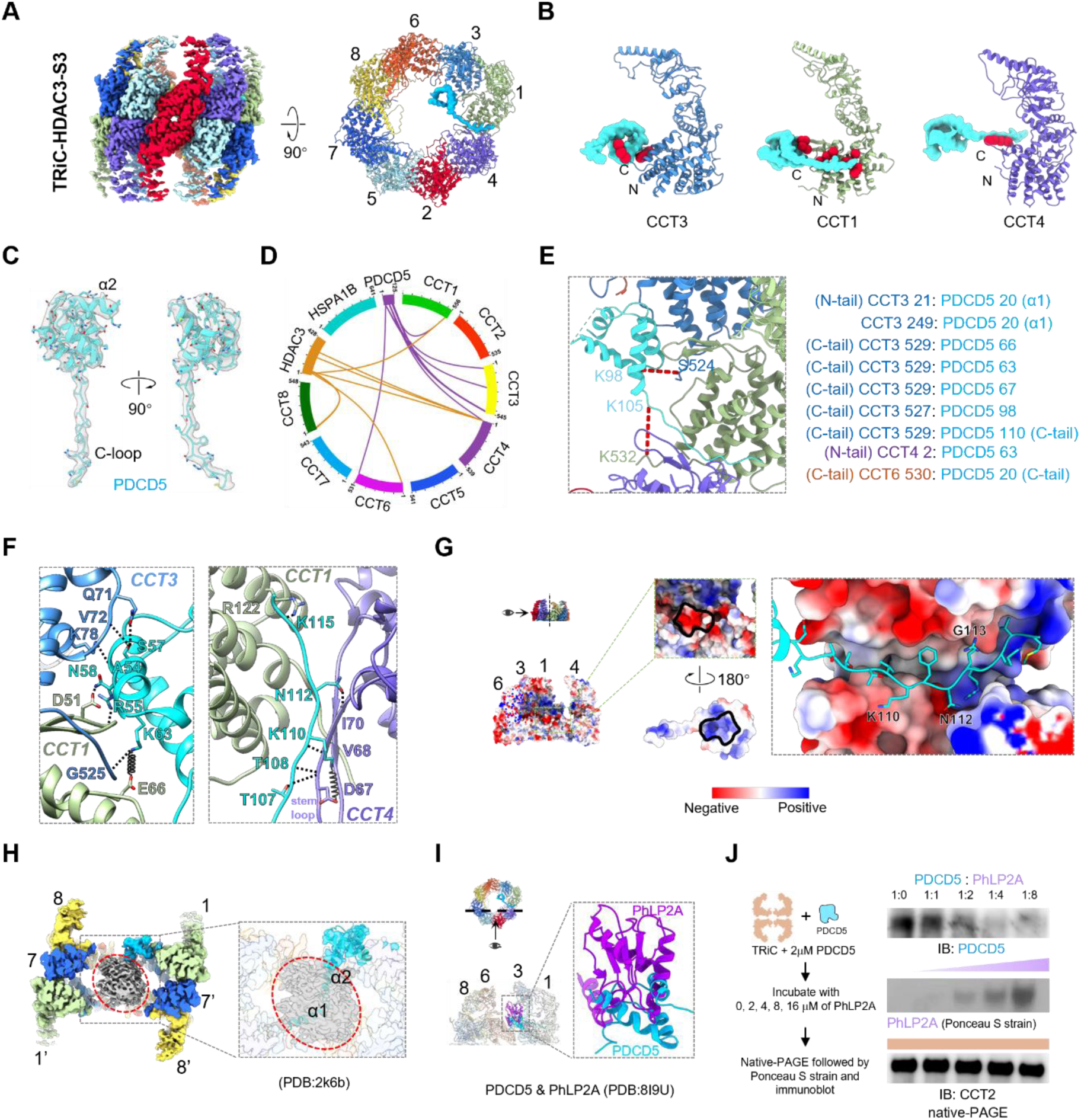
Cryo-EM structures of HDAC3-TRiC in the open S3 state. (A) Side and top views of the cryo-EM maps of HDAC3-TRiC in the S3 states, revealing one or two extra densities (aquamarine) within the TRiC chamber. (B) TRiC-PDCD5 interaction analysis, showing PDCD5 contacts with CCT3/1/4 subunits. TRiC residues within 4 Å of PDCD5 are shown as red balls. (C) The extra density within the TRiC chamber closely matches the AlphaFold2-predicted PDCD5 model. (D) XL-MS analysis of the HDAC3-TRiC complex identifies cross-links between HDAC3, PDCD5, and TRiC. (E) Intermolecular PDCD5-TRiC crosslinks mapped onto the model, with cross-links to disordered N-/C-terminal tails listed on the left. (F) Interaction networks between PDCD5 and TRiC. (G) Electrostatic surface property analyses depict PDCD5 engages TRiC via electrostatic interactions. (H) Clipped view of the unsharpened S3 map, highlighting PDCD5 densities (transparent aquamarine) contacting TRiC’s unstructured N-/C-terminal tails (grey, within the red circle). The NMR-resolved α1-helix of PDCD5 (pdb:2K6B) may clash with these tails. (I) Overlay of our PDCD5-TRiC and the PhLP2A-TRiC models (PhLP2A in purple), showing steric clashes between PDCD5 and PhLP2A. (J) PDCD5-TRiC binding competition with PhLP2A. PDCD5 was incubated with TRiC on ice, after that different concentrations of PhLP2A were added. The samples were analyzed via native-PAGE, followed by Ponceau S staining and immunoblotting.

Our map resolves most of PDCD5, except the α1- and part of the α2-helix (Fig. 4C, S5F). Its α3-helix interacts with the CCT1 stem-loop and CCT3 H3-helix primarily via H-bonds, while its C-terminal tail occupies the CCT1-CCT4 E-domain cleft, forming extensive H-bonds and salt bridges, especially with the CCT4 stem-loop (Fig. 4B, F, Table S5). TRiC and PDCD5 interact via electrostatic interactions (Fig. 4G). Moreover, based on its α2-helix orientation contacting TRiC’s central density, which corresponds to disordered N- and C-terminal tails^4,44,45^, and the NMR structure of free PDCD5 ^50^, its α1-helix could extend into this tail region (Fig. 4H). XL-MS revealed numerous crosslinks between PDCD5 and TRiC’s terminal tails (Fig. 4D-E), confirming interaction with TRiC E-domain and disordered terminal regions. Corroborate our structural observations, no Hsp70-TRiC crosslinks were detected in the XL-MS data (Fig. 4D), indicating Hsp70 dissociates from TRiC upon nucleotide binding.

Prior studies reported that PDCD5 interacts with HDAC3 ^51^, inhibiting HDAC3 function in certain cells^51,52^. We propose that PDCD5 associates with HDAC3 via TRiC, suppressing HDAC3 function by inhibiting its folding. Furthermore, PDCD5 engages the CCT1/CCT4 stem-loops and TRiC’s N- and C-terminal regions (Fig. 4F-H)—key allosteric elements mediating TRiC ring closure^15,44^. Thus, PDCD5 may regulate TRiC ring closure, potentially suppressing TRiC-mediated HDAC3 folding.

### Folding of other class I HDACs: HDAC1 requires TRiC, but HDAC8 does not

Our study elucidates TRiC-assisted HDAC3 folding, orchestrated by a co-chaperone and cofactor network including Hsp70 and PDCD5. Given the high sequence identity among class I HDACs (59.1% between HDAC3 and HDAC1/2; 41% between HDAC3 and HDAC8; Fig. 5A, S6A-B)^32^, we hypothesized that TRiC broadly assist their folding. Co-IP confirmed TRiC-HDAC1 interactions (Fig. 5B), aligns with a prior report^53^. To validate this, we overexpressed Flag-tagged HDAC1 in HEK293F cells, and affinity purification followed by glycerol density gradient centrifugation indeed isolated a TRiC-HDAC1 complex (Fig. S7C), verifying their direct interaction.

**Fig. 5.**
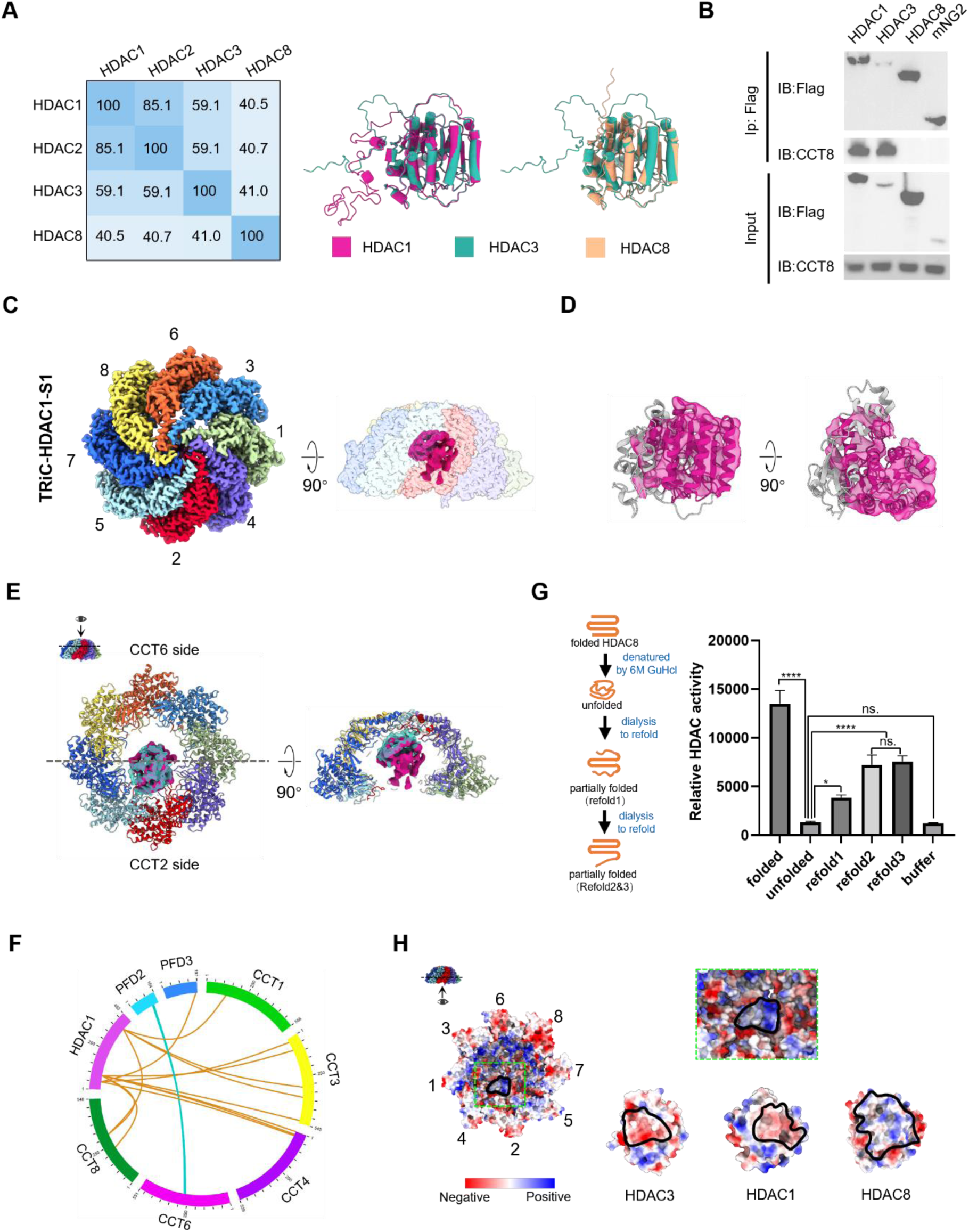
Cryo-EM structure of closed HDAC1-TRiC-S1 reveals TRiC-assisted folding of HDAC1. (A) Sequence identities of HDAC1, HDAC2, HDAC3, and HDAC8. AlphaFold2-predicted HDAC3 model were aligned to HDAC1 and HDAC8. (B) Co-immunoprecipitation (co-IP) detected interactions between HDAC1/HDAC3/HDAC8 and TRiC. (C) Top and side views of the cryo-EM map of HDAC1-TRiC in the closed S1 state, exhibiting extra densities (medium violet red) within TRiC chamber. (D) The extra density within TRiC chamber closely matches the HDAC1 model, with the N-terminus slightly less well folded. (E) Comparison of substrate density locations in HDAC1-TRiC-S1 and HDAC3-TRiC-S2, with their structures superimposed (HDAC3 density transparent seagrass green, HDAC3 violet red). (F) XL-MS analysis of the HDAC1-TRiC complex, showing cross-links between HDAC1, TRiC, and PFD. (G) HDAC8, denatured with 6M guanidine hydrochloride and dialyzed to remove denaturing agent, was monitored for enzymatic activity recovery during dialysis. (H) Electrostatic surface property analysis reveals that, when aligned with HDAC1 and HDAC3 in the TRiC chamber, HDAC8’s potential TRiC-interaction surface is predominantly positively charged, opposite those of HDAC1 and HDAC3, disfavoring TRiC binding.

Cryo-EM analysis of the TRiC-HDAC1 complex in the presence of ATP-AlFx revealed three conformational states—TRiC-HDAC1-S1, -S2, and -S3, at 3.64 Å, 3.89 Å, and 3.55 Å resolution, respectively (Fig. S7A-B). The TRiC-HDAC1-S1 map shows TRiC adopts a closed conformation featuring an additional internal density matching HDAC1 well (Fig. 5C-D, S7C-D). Notably, the shape, orientation, and binding position of HDAC1 mirror those of HDAC3 (Fig. 5E), suggesting a conserved TRiC-mediated folding mechanism for both HDAC1 and HDAC3. XL-MS confirmed direct HDAC1-TRiC interactions (Fig. 5F, Table S6), aligning with our structural and biochemical findings.

In contrast, despite 41% sequence identity with HDAC1/HDAC3 (Fig. 5A, S6B), HDAC8 showed no interaction with TRiC in Co-IP experiment (Fig. 5B). Overexpression of Flag-tagged HDAC8 in HEK293F cells, followed by affinity purification, yielded pure HDAC8 without TRiC or other chaperones (Fig. S6D). Unlike HDAC1/3, HDAC8 retains enzymatic activity when expressed in *E. coli* ^54,55^. These findings suggest that HDAC8 folding bypasses TRiC. To confirm this, we denatured purified HDAC8 with 6M guanidine hydrochloride (GuHCl) and gradually removed the denaturant via dialysis. Enzymatic activity assays revealed partial activity recovery (Fig. 5G), verifying that HDAC8 folds independently of TRiC.

In summary, our findings establish that TRiC is essential for folding HDAC1 and HDAC3 (and likely HDAC2, given its 85.1% identity to HDAC1), but not HDAC8, the only class I HDAC functioning as an isolated enzyme^34,35^. Modeling HDAC8 in an orientation similar to HDAC1 and HDAC3 reveals a predominantly positively charged potential TRiC-binding surface, in contrast the favorable interfaces of HDAC1 and HDAC3 (Fig. 5H), likely preventing TRiC interaction. These results underscore the sophisticated substrate specificity of TRiC and its co-chaperones among class I HDACs, where subtle structural variations govern chaperone dependence and folding process.

### HDAC1 employs PFD to orchestrate its TRiC-assisted folding

We demonstrated that Hsp70 and PDCD5 mediate TRiC-assisted HDAC3 folding. To investigate whether these factors also contribute to HDAC1 folding, we analyzed open TRiC particles from the TRiC-HDAC1-ATP-AlFx dataset, resolving two conformational states, designated TRiC-HDAC1-S2 and TRiC-HDAC1-S3 (Fig. S7A-B). Strikingly, the S2 map revealed a PFD complex bound atop TRiC, engaging the apical protrusions of CCT3 and CCT4 (Fig. 6A, S8A-C). Specifically, the CCT4 H8-helix inserts between PFD4 and PFD6 subunits (Fig. 6A), consistent with previous observations in TRiC-tubulin system^13,14^. Overlaying the TRiC-HDAC1-S2 and TRiC-HDAC3-S1 maps showed that PFD and Hsp70 target overlapping CCT4 A-domain sites (Fig. 6B), suggesting competitive, mutually exclusive binding. Thus, despite both class I HDACs, HDAC1 and HDAC3 rely on distinct substrate-delivery co-chaperones—PFD for HDAC1 and Hsp70 for HDAC3—to facilitate their TRiC-mediated folding.

**Fig. 6.**
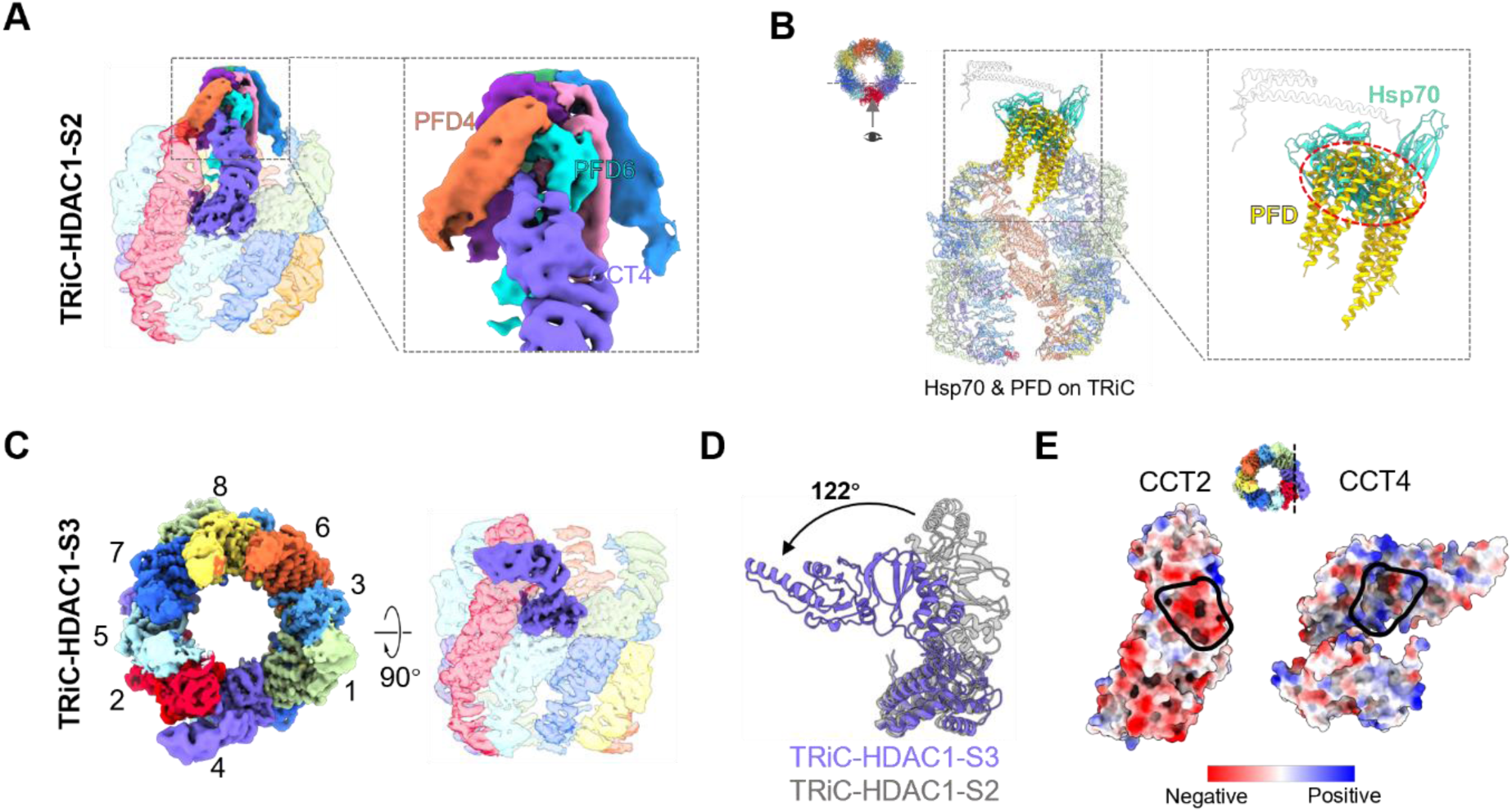
Cryo-EM structures of the open HDAC1-TRiC-S2 and -S3 states reveal PFD binding to and dissociation from TRiC. (A) Cryo-EM map of the HDAC1-TRiC-S2, showing PFD bound atop TRiC (transparent). CCT4 apical protruding H8-helix inserts between PFD4 and PFD6. (B) Steric clashes between Hsp70 and PDF atop TRiC, by overlaiding HDAC1-TRiC-S2 and HDAC3-TRiC-S1. (C) Cryo-EM map of the HDAC1-TRiC-S3 state. (D) In the S3 state, CCT4 exhibits a 122°A- and I-domain bend compared to its typical upright conformation in S2 state. (E) Electrostatic surface property analyses suggest that the bent CCT4 interacts with CCT2 through electrostatic interaction.

### Dramatic CCT4 A- and I-domain bending may regulate co-chaperone release

Unexpectedly, the TRiC-HDAC1-S3 open-state map captures an unprecedented TRiC conformation (Fig. 6C, S8D-F), characterized by a dramatic 122°counterclockwise rotation of CCT4’s A- and I-domains (Fig. 6D), resembling CCT2’s bent conformation in yeast TRiC in NPP state^45^ (Fig. S8G). This CCT4 repositioning establishes electrostatic interactions with the back of CCT2 I-domain (Fig. 6E). Notably, from TRiC-HDAC1-S2 to S3 state, this pronounced rearrangement retracts CCT4’s H8-helix from PFD4/6 subunits, likely facilitating PFD dissociation from TRiC post-substrate delivery. These dindings suggest that CCT4’s A- and I-domain bending orchestrates PFD binding and release from TRiC (Fig. 6C). Similarly, our TRiC-HDAC3-S1 map shows Hsp70 binding to the CCT4 A-domain (Fig. 2E), implying that CCT4 bending may also trigger Hsp70 release. Specifically, in its upright conformation, CCT4 engages PFD or Hsp70 to facilitate substrate transfer; bending likely induces their dissociation from TRiC. Thus, the conformational plasticity of CCT4 acts as a molecular switch coordinating co-chaperone binding and release.

Intriguingly, despite using identical endogenous substrate affinity purification strategy, PDCD5 was exclusively detected in the TRiC-HDAC3 complex, not in TRiC-HDAC1, suggesting substrate-specific PDCD5 interactions.

## Discussion

Class I HDACs—HDAC1, HDAC2, HDAC3, and HDAC8—are nuclear enzymes critical for regulating cell survival, differentiation, proliferation, and apoptosis^26^. Using an integrated approach combing cryo-EM, biochemical assays, and XL-MS, we elucidated the unexpected TRiC-assisted folding pathways and mechanisms of HDACs, in concert with a dynamic co-chaperone/cofactor network. For HDAC3, our findings suggest that Hsp70, positioned atop TRiC (Fig. 2A-G), likely facilitates HDAC3 delivery, while PDCD5, located within the open TRiC chamber (Fig. 4A), may modulate its folding. We identified HDAC3’s H14 helix as a critical structural determinant in this process (Fig. 3H). Extending beyond HDAC3, we established that HDAC1 (and likely HDAC2, given its 85.1% sequence identity to HDAC1) is also a TRiC substrate and resolved the open and closed TRiC-HDAC1 structures. Strikingly, HDAC1 employs another co-chaperone, PFD, for delivery to TRiC (Fig. 6A), yet shares core TRiC-assisted folding features with HDAC3. In contrast, HDAC8—uniquely active as an isolated enzyme in class I HDACs^34,35^— folds independently of TRiC (Fig. 5B, G), revealing a profound divergence in chaperonine and co-chaperone dependency among class I HDACs. Collectively, our results provide a complete picture of the TRiC-mediated folding process and mechanisms of class I HDAC family members, in precise coordination with a dynamic network of co-chaperones and cofactors, and uncover emerging working principles governing the regulation of TRiC by these co-chaperones and partner cofactors.

TRiC facilitates the folding of a highly diverse array of substrates. Here, we showed that its functional diversity is further enhanced by intricate interplay with co-chaperone/cofactor networks, which add an additional layer of regulation on substrate selection, delivery, folding, and release^1^ (Fig. 7A-B). We elucidate the mechanisms of TRiC-assisted folding of class I HDACs, revealing a finely tuned substrate selection process mediated by co-chaperones and cofactors. Specifically, HDAC1 and HDAC3 rely on TRiC for folding but employ distinct co-chaperone/cofactor: Hsp70 and PDCD5 regulate HDAC3 delivery and folding, whereas PFD mediates HDAC1 delivery. In contract, HDAC8—despite sharing 41% sequence identity and structural similarity with HDAC1/3—folds independently of TRiC. Structural analyses suggest that subtle differences in sequence or surface properties between HDAC1/3 and HDAC8 (Fig. 5H) may contribute to their divergent dependence on TRiC and its co-chaperone/cofactor networks.

**Fig. 7.**
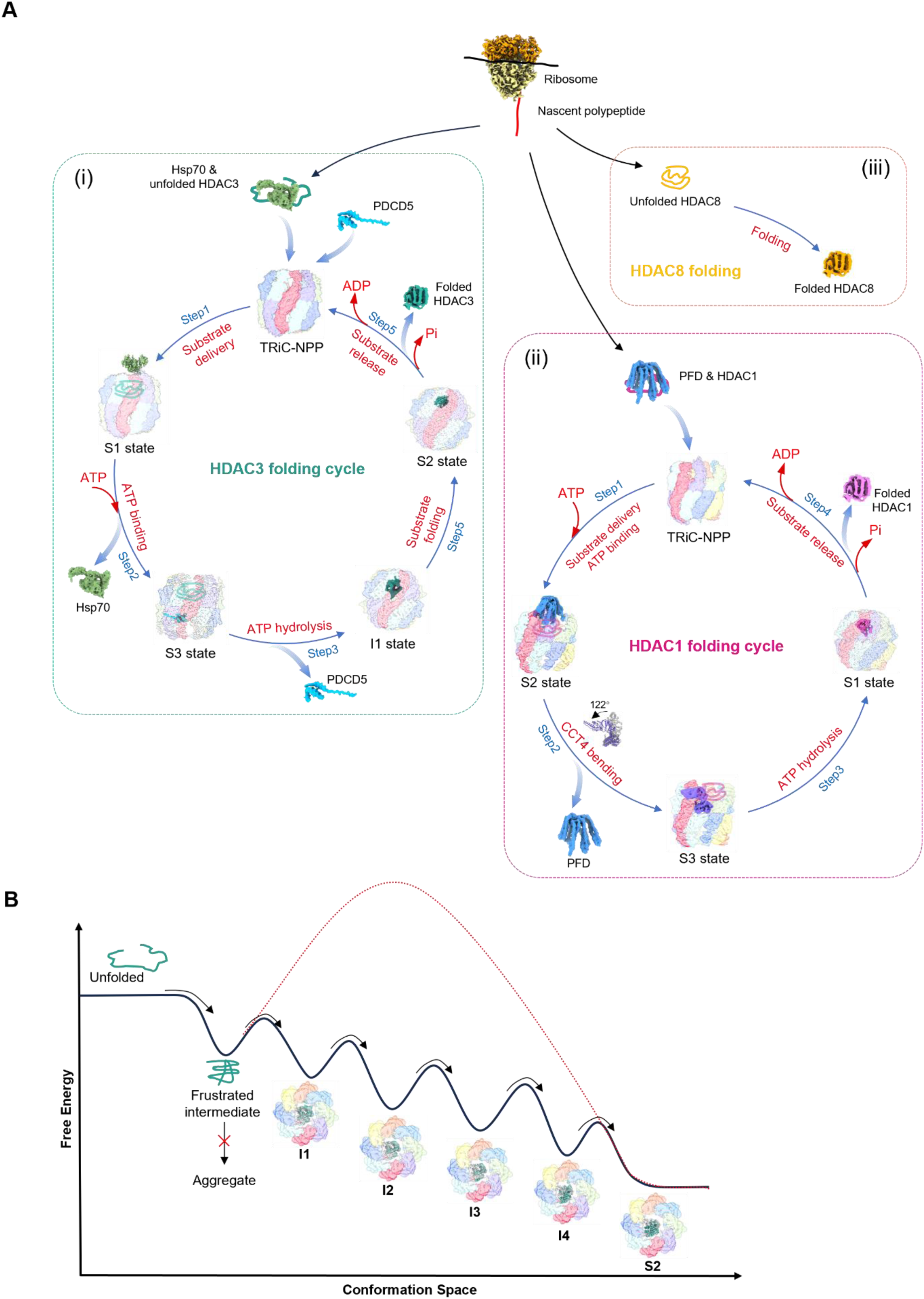
Pathway and mechanism of TRiC-mediated folding of Class I HDACs in coordination with co-chapernone/cofactor networks. (A) Folding cycle of Class I HDACs. (i) After ribosomal translation, unfolded HDAC3 binds Hsp70, which lands atop the open NPP-state TRiC, delivering HDAC3 into the TRiC chamber (step 1). ATP binding triggers Hsp70 dissociation, stabilizing PDCD5 within the chamber, potentially regulating TRiC ring closure (step 2). ATP hydrolysis then drives TRiC ring closure, releasing PDCD5 and initiating HDAC3 folding. The Η14-helix emerges first, followed by folding propagation to nearby elements and eventually to the remaining regions. This illustrates a stepsie folding pathway, with the folded HDAC3 positioned on the CCT2 side of TRiC (step 3). Upon Pi release and TRiC chamber reopening, folded HDAC3 is released and assembles into a functional complex (step 4). (ii) For HDAC1, post-translation, PFD could associate and deliver it to the TRiC chamber (step 1). Bending of CCT4’s A- and I-domains induces PFD release (step 2). ATP hydrolysis drives TRiC ring closure, facilitating HDAC1 folding (step 3). Following Pi release and TRiC chamber reopening, folded HDAC1 is released and assembled into a functional complex (step 4). (iii) In contrast, HDAC8 likely folds spontaneously post-translation, functioning independently of TRiC and co-chaperone/cofactors. (B) Proposed energy landscape of TRiC-assisted HDAC3 folding. Newly synthesized HDAC3, in the absence of TRiC chaperonin assistance, is prone to aggregation or must overcome a substantial energy barrier to attain its native conformation. Engagement of HDAC3 with TRiC, coupled with its ATPase cycle, likely restricts the conformational landscape of HDAC3 and reduces its folding energy barrier. This chaperone-mediated confinement stabilizes HDAC3 intermediates, enabling progressive folding into a functional near-native state.

Our results indicate that for TRiC, its parallel functional co-chaperones or cofactors may operate in a mutually exclusive manner, reflecting a sophisticated regulatory framework. Specifically, Hsp70 and PFD, which deliver HDAC3 and HDAC1/tubulin/actin, respectively, compete for an overlapping CCT4 A-domain binding site atop TRiC, precluding simultaneous binding (Fig. 6B). Similarly, cofactors PDCD5 and PhLP2A, coordinating with HDAC3 and tubulin/actin^15,17^, respectively, target overlapping regions on CCT3 subunit within the open TRiC chamber (Fig. 4I). Indeed, native-PAGE assays confirm this exclusivity, showing PhLP2A competes with PDCD5 for TRiC binding (Fig. 4J). A recent *in situ* cryo-ET study suggested PDCD5 associates with nearly all open TRiC complexes in HEK293F cells, and reported simultaneous PDCD5 and PFD co-binding^19^. Still, in our class I HDAC-overexpressed system, structural analyses detect no such co-occupancy. Together, these findings highlight the intricate spatial and temporal regulation and substrate specificity governing the interplay between TRiC and its co-chaperones and cofactors, ensuring precise chaperonin function.

Furthermore, our findings suggest that TRiC may regulate co-chaperone recruitment and release through CCT4 conformational changes. In its upright conformation, CCT4 accommodates PFD binding atop TRiC, whereas its transition to a bent conformation promotes PFD dissociation. A parallel mechanism may apply to Hsp70, as both co-chaperones primarily target the CCT4 A-domain’s apical protrusion region. Additionally, the CCT4 E-domain recruits cofactors PDCD5 and PhLP2A^17^ within the TRiC chamber. Notably, lethal mutations in CCT4’s nucleotide-binding pocket in yeast^56^ suggest that disrupted ATPase activity may impaired CCT4’s coordination of co-chaperone and cofactor binding and release. Collectively, these observations position CCT4 conformational changes as a key mechanism coordinating TRiC’s dynamic interplay with its co-chaperone/cofactor network.

The special and temporal localization of TRiC and its substrates may also impact cellar processes. HDAC3, primarily localized in the nucleus, forms complexes with NCOR1 and SMRT^33^, while HDAC1, also predominantly nucleus, forms complexes such as Sin3A^30^ and NuRD^28^. Their enzymatic activities are tightly modulated by interacting proteins^32,35,37^. Notably, ∼90% of TRiC substrates are components of oligomeric complexes, relying on TRiC not only for folding but also potentially for assembly^2,57^. Protein translation occurs in the cytoplasmic matrix, and their folding is tightly regulated. Although TRiC is primarily cytoplasmic, we and others have detected it in the nucleus (Fig. S10) ^58,59^. This dual localization raises the possibility that TRiC may engage newly translated HDAC1 and HDAC3 in the cytoplasm, facilitating their folding followed by transport into the nucleus, where it could further contribute to their assembly into functional complexes. Additionally, TRiC interacts with components of nuclear complexes, including SAGA^60^, SWI/SNF^61^, Set1/compass^61^, CRL^CSA^ DNA-repair factor^62^, TFIID^60^, and RNA polymerase II^61^, indicating a role in transcriptional regulation by contributing to the folding and assembly of these machineries. Collectively, these findings underscore TRiC’s versatile roles in cellular proteostasis across compartmental boundaries.

In summary, this study elucidates the mechanisms governing TRiC-assisted folding of class I HDACs, revealing distinct co-chaperone and cofactor dependencies for HDAC1 and HDAC3, whereas HDAC8 folds independently of TRiC. Furthermore, we uncover that TRiC-co-chaperone/cofactor interactions are modulated by conformational changes in the CCT4 subunit. Our results reveal that subtle structural variations among substrates, coupled with their spatiotemporal localization, influence TRiC-mediated substrate folding and co-chaperone/cofactor specificity. These findings highlight the sophisticated regulatory landscape of TRiC and open promising avenues for designing peptides or small molecules to selectively modulate TRiC-assisted folding of class I HDACs and other substrates.

